# Convergent effects of different anesthetics on changes in phase alignment of cortical oscillations

**DOI:** 10.1101/2024.03.20.585943

**Authors:** Alexandra G. Bardon, Jesus J. Ballesteros, Scott L. Brincat, Jefferson E. Roy, Meredith K. Mahnke, Yumiko Ishizawa, Emery N. Brown, Earl K. Miller

## Abstract

Many anesthetics cause loss of responsiveness despite having diverse underlying molecular and circuit actions. To explore the convergent effects of these drugs, we examined how anesthetic doses of ketamine and dexmedetomidine affected oscillations in the prefrontal cortex of nonhuman primates. Both anesthetics caused increases in phase locking in the ventrolateral and dorsolateral prefrontal cortex, within and across hemispheres. However, the nature of the phase locking varied. Activity in different subregions within a hemisphere became more anti-phase with both drugs. Local analyses within a region suggested that this finding could be explained by broad cortical distance-based effects, such as large traveling waves. By contrast, homologous areas across hemispheres became more in-phase. Our results suggest that both anesthetics induce strong patterns of cortical phase alignment that are markedly different from those in the awake state, and that these patterns may be a common feature driving loss of responsiveness from different anesthetic drugs.

## Introduction

Many different anesthetics exist and are widely used in medicine for similar purposes, despite having distinct mechanisms of action. The common elements by which these drugs induce similar outcomes, namely loss of responsiveness, is an open question^1,2^. Their convergence is likely at a higher level, in the network dynamics driven by each drug’s unique molecular effects. Here, we explored the idea that different anesthetics may cause loss of responsiveness via similar effects on cortical synchrony.

To address the question of functional convergence we examined ketamine and dexmedetomidine, two anesthetics with distinct molecular actions on different pathways. Ketamine strongly affects the cortex by antagonizing N-methyl-D-aspartate (NMDA) receptors. By contrast, dexmedetomidine agonizes α2 adrenergic receptors, mainly in the locus coeruleus. At anesthetic doses, they both produce the same behavioral outcome: loss of responsiveness.

We examined bilateral recordings from the prefrontal cortex (PFC) of macaques during administration of ketamine and dexmedetomidine. We quantified cortical synchrony, which is thought to play a role in consciousness^3–8^. We explored not only phase locking, a common measure of the consistency of local field potential (LFP) signals between different areas, but also the phase offsets at which these LFPs were locked.

We found evidence that ketamine and dexmedetomidine had similar effects in how they changed phase relationships between low-frequency LFPs. The drugs increased phase locking across all recorded regions of the prefrontal cortex, but the nature of this phase locking was not homogeneous. Between different regions within a hemisphere, the drugs both induced consistently misaligned LFP activity, while between homologous regions across hemispheres, they caused activity to become more aligned. These shifts suggest that anesthetics act through complex changes in phase alignment of oscillatory dynamics that are common across drugs despite their different molecular actions.

## Results

We examined activity simultaneously recorded from electrode arrays in the right dorsolateral and ventrolateral PFCs (R-Dor and R-Ven, respectively) and left dorsolateral and ventrolateral PFCs (L-Dor and L-Ven, respectively). A lever-pressing task, which involved pressing a lever for a reward whenever a tone was played, was used to assess loss of responsiveness. The task was run before anesthetic drug administration, around the time of administration, and during the period of unresponsiveness induced by the anesthetics. The lever pressing task occurred during periods with a white background in Figure 1A. Anesthetic doses of ketamine (10 mg/kg) and dexmedetomidine (20 mg/kg) caused monkeys to stop responding (dots indicate responses, Fig. 1A) shortly after their administration (vertical line at time zero, Fig. 1A), indicating the onset of anesthetic effects. For all the following analyses, neural data was taken from periods during which the lever-pressing task was not happening (gray boxes, Fig. 1A), to minimize differences between before- and during-anesthetic periods due to movement and reward. Animals remained unresponsive even after the lever task was reinstated at 25 min post drug. LFPs were pre-processed using a statistically robust common average referencing technique (rCAR), which has been shown to more accurately maintain a signal’s phase than other methods^9^ (see “LFP Processing” section of Methods for a discussion of referencing techniques).

**Figure 1:**
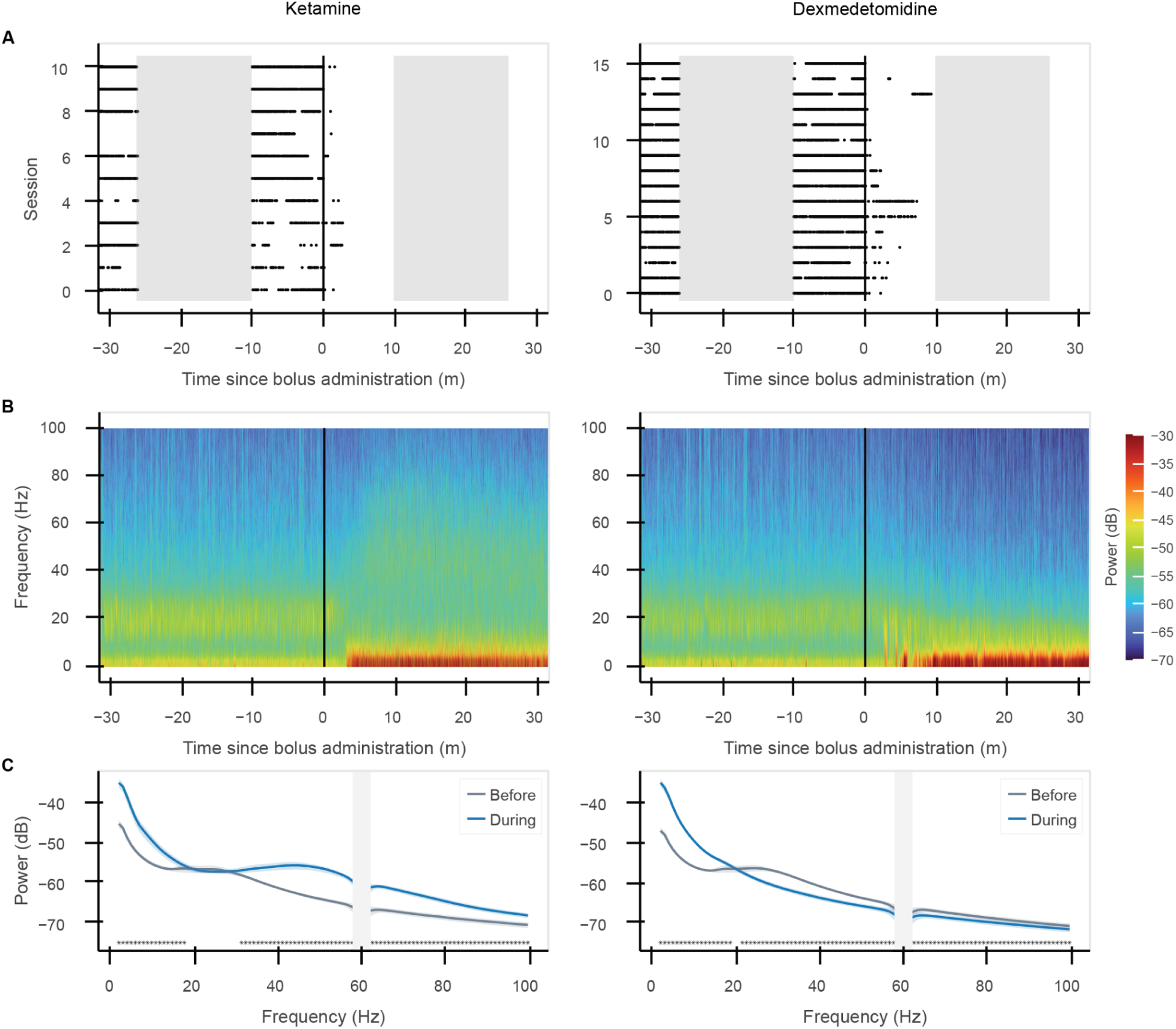
Both anesthetics induced loss of responsiveness, increased low-frequency power. (A) Behavioral response to the lever-pressing task. Black dots indicate lever presses; black line at time 0 indicates drug administration. Gray areas indicate time periods without the lever task (further analysis focuses on data from this period). (B) LFP power as a function of frequency and time for two representative sessions (single channel), one with ketamine (left) and one with dexmedetomidine (right). Black line at time 0 indicates drug administration. (C) Average LFP power across all sessions before (gray) and during (blue) anesthetics (mean and 99% confidence intervals across sessions). Stars indicate significant differences in power before and during anesthetic effects (p&lt;0.01, corrected for multiple comparisons). Gray bar at 60 Hz due to line noise filtering effects.

## Anesthetics increased low-frequency power

Ketamine and dexmedetomidine at anesthetic doses increased low-frequency LFP power. Figure 1B shows the spectrogram of LFP power from one channel of one array around the administration of anesthetics for a representative session from each drug type. Across all sessions, both drugs caused an increase in power below 16 Hz that was especially prominent in the delta (1-4 Hz) and theta (4-8 Hz) bands (Fig. 1C). Ketamine also caused an increase in gamma band power (30-100 Hz), while dexmedetomidine caused a slight decrease (Fig. 1C).

## Anesthetics increased LFP phase locking

Both drugs caused an increase in phase locking at low frequencies. Figure 2A shows phase locking, as measured by the phase-locking value (PLV), before drug delivery. It shows average phase locking between LFPs from all possible pairs of electrodes in the four arrays implanted in the left and right dorsolateral and ventrolateral PFC. Here and elsewhere, comparisons within hemispheres are colored green and those across hemispheres are colored purple (Fig. 2).

**Figure 2:**
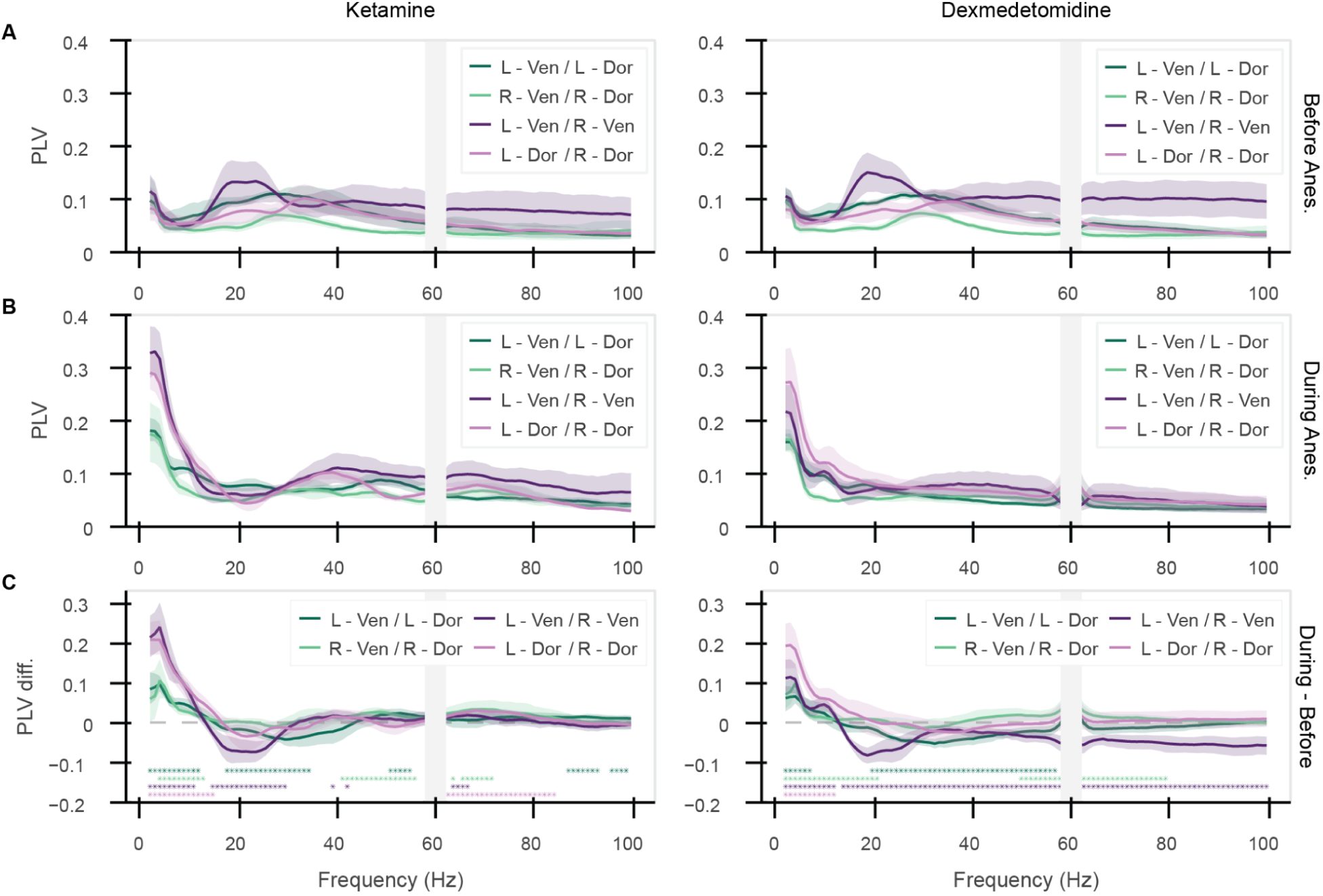
Anesthetics increased LFP phase locking. Phase-locking value (PLV) between channels in pairs of arrays (mean and 99% confidence intervals across sessions). (A) Before anesthetics, (B) during anesthetics, (C) their difference (during – before). Stars indicate significant differences in phase locking before and during anesthetic effects (p&lt;0.01, corrected for multiple comparisons). Gray bar at 60 Hz due to line noise filtering effects.

After drug delivery, there was a significant increase in average phase locking at low frequencies (1-8 Hz) between the dorsolateral and ventrolateral PFC within each hemisphere, as well as across the two hemispheres (Fig. 2B). Figure 2C shows the change in phase locking caused by the drugs. The increase indicated that the relative phase between pairs of LFPs was becoming more consistent. Note that this increase in phase locking did not necessarily mean the phases were aligned, a point we will take up below. There was a dip in phase locking between ventrolateral PFC in either hemisphere in the beta range (13-30 Hz), as well as in the gamma range for dexmedetomidine (60-100 Hz), due to high phase locking in this range before anesthetics were administered. There were additional significant changes in phase locking at higher frequencies, but these effects were smaller than those seen at the low frequencies.

We confirmed this effect using another measure of phase locking, coherence. PLV measures phase locking independently of changes in power^10–12^. Coherence takes into account both phase and power to measure how linearly predictable one signal is from another. This analysis yielded similar results: an increase in signal consistency at low frequencies (Fig. S1A). The effects were similar for all combinations of the arrays, both within hemispheres (R-Dor vs. R-Ven, L-Dor vs. L-Ven) and across hemispheres (R-Dor vs. L-Dor and R-Ven vs. L-Ven) (Fig. S1A). There were also similar trends for the “diagonal” comparisons across both subregions *and* hemispheres (R-Dor vs. L-Ven and L-Dor vs. R-Ven), though these effects did not always reach significance due to increased variability (Fig. S1B-C). While there was variation between the two monkeys, likely due to a combination of biological variation and slight differences in array placement, these effects in the low frequencies were generally directionally consistent across different sessions and across the two monkeys (Fig. S2A-B).

## Anesthetics misaligned activity within hemispheres but aligned it across hemispheres

The increase in low-frequency phase locking caused by ketamine and dexmedetomidine at anesthetic doses did not necessarily mean that the phases became more aligned. We found that the phase offsets became more anti-phase within hemispheres and more in-phase across hemispheres.

Figure 3A and B show the distribution of phase offsets in the 1-4 Hz range. The gray bars show the phase offsets before drug delivery. They were nearly uniform across hemispheres and biased toward in-phase (0° offset) within hemispheres. The colored bars show phase offsets after anesthetic administration. They indicate an increase in anti-phase offsets between ventrolateral and dorsolateral arrays (L-Ven vs. L-Dor, R-Ven vs. R-Dor; Fig. 3A-B, green). By contrast, there was increased phase alignment across hemispheres (L-Ven vs. R-Ven, L-Dor vs. R-Dor; Fig. 3A-B, purple).

**Figure 3:**
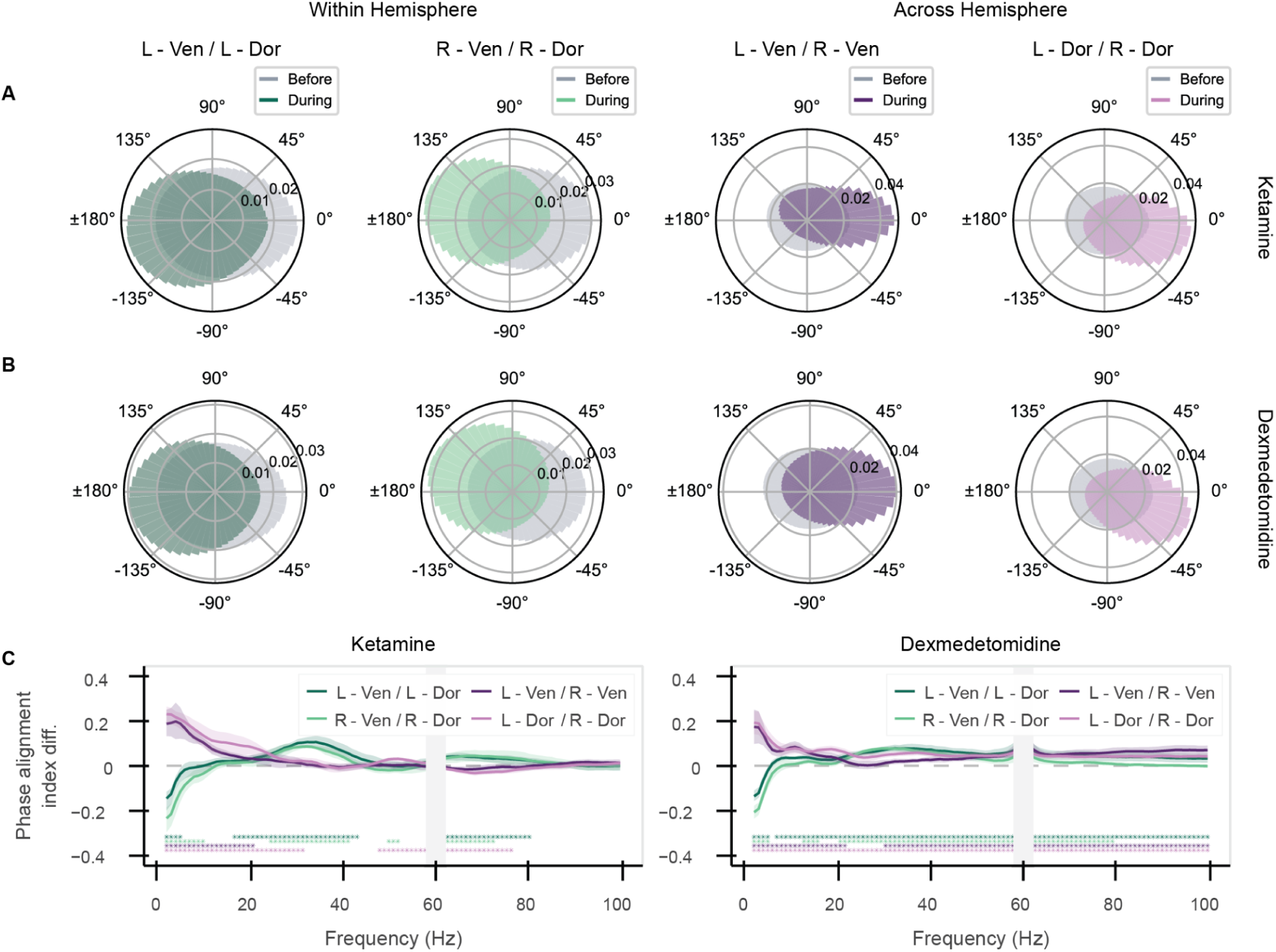
Anesthetics increased phase offsets within hemispheres, decreased across hemispheres. (A-B) Histogram of phase offsets between all channel pairs in indicated regions before (gray) and during (colored) anesthetics for (A) ketamine and (B) dexmedetomidine in the 1-4 Hz range. (C) Phase alignment index cos(Δϕ) difference (during – before anesthetics) (mean and 99% confidence intervals across sessions). 0° offset is +1, 180° offset is -1, and uniform offset averages to zero. Stars indicate significant differences in phase alignment before and during anesthetic effects (p&lt;0.01, corrected for multiple comparisons). Gray bar at 60 Hz due to line noise filtering effects.

The changes in phase offsets were mainly in the lower frequencies. To quantify this effect, we calculated a phase alignment index, measuring the average cosine of the phase offsets. Figure 3C shows the change in the phase alignment index with anesthetics (non-subtracted values before and during anesthetics shown in Fig. S3). Anesthetic doses of both ketamine and dexmedetomidine caused a significant decrease in phase alignment within hemispheres (negative alignment index, more towards 180°) at frequencies in the 1-4 Hz range (Fig. 3C, green lines). Across hemispheres, there was a significant increase in phase alignment (positive alignment index, more towards 0°) for the same frequency range (Fig. 3C, purple lines).

Ketamine also caused a significant increase in within-hemisphere phase alignment in the 20-40 Hz range, and both drugs had a range of small but statistically significant effects on phase alignment in the higher frequencies. The effects were directionally consistent across animals, sessions, and drugs (Fig. S4). Along the diagonal comparisons (L-Ven vs. R-Dor, R-Ven vs. L-Dor) we also found a slight increase in anti-phase signals with dexmedetomidine, but not ketamine (Fig. S5). In addition, we tested the changes in phase offsets with multiple LFP referencing techniques to ensure that our findings did not rely upon the referencing scheme. In addition to the method used here, rCAR, we tested no re-referencing and global mean subtraction (see Methods for further discussion of referencing). All techniques led to similar trends in changes in phase offsets with anesthetics (Fig. S6).

## Increased phase offset with distance was amplified by anesthetics

To explore whether the observed cross-region phase offsets could be explained by distance effects, we quantified how the phase locking and phase offsets changed with distance between electrodes within an array, where precise relative locations of recording sites were known. We found that in the awake PFC, phase locking decreased and phase offset increased with distance within each recording array. These effects were stronger with anesthetics. The phase locking and phase offsets were relatively linear as a function of distance (example of low-frequency PLV in Fig. 4A), so we also examined the slopes of these lines.

**Figure 4:**
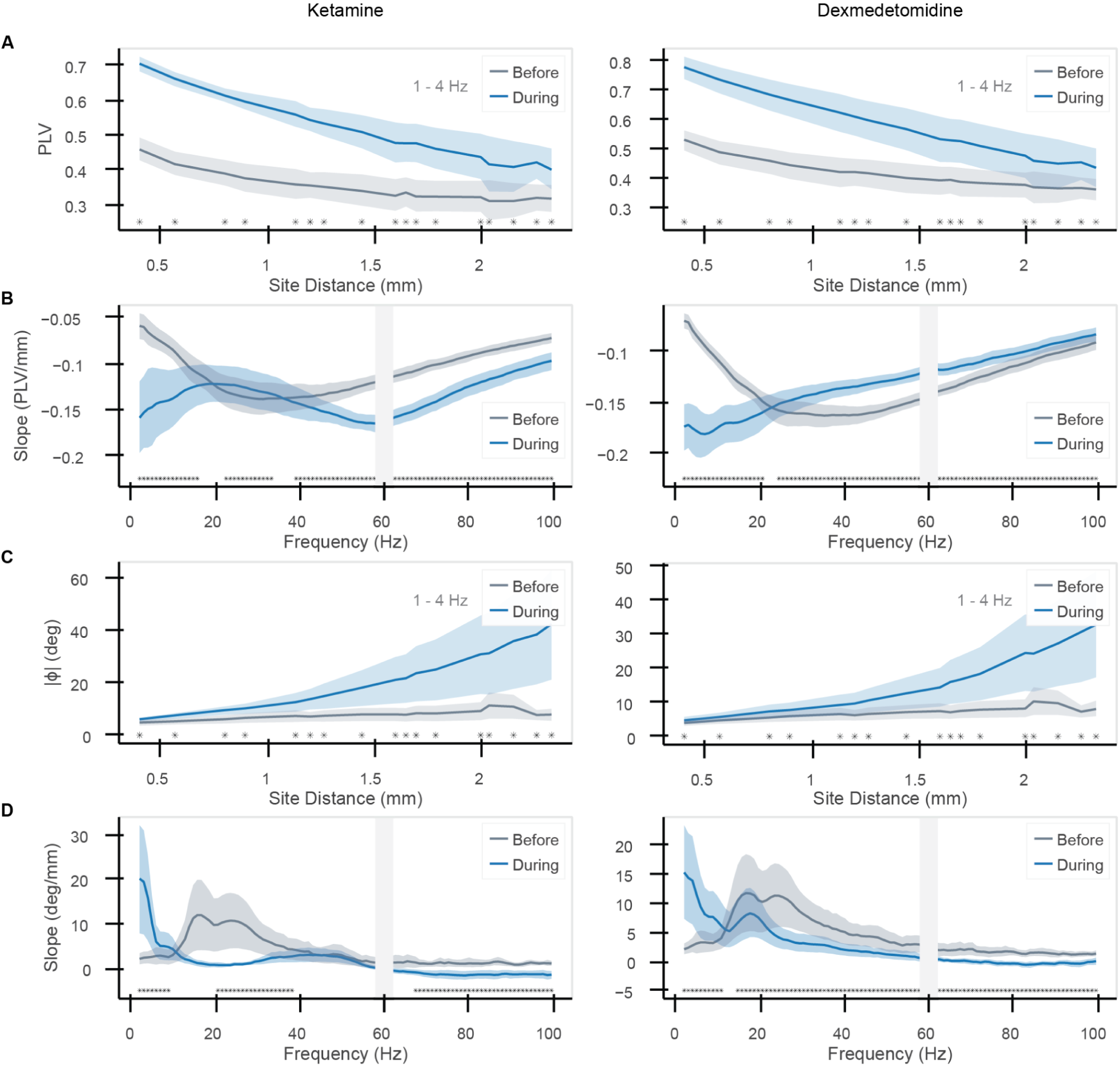
Phase locking weakened and LFPs became more anti-phase with distance. (A) PLV vs. distance within an array, before (gray) and during (blue) anesthetics, in the low-frequency range (1-4 Hz). (B) Slope of PLV vs. distance (as in (A)) as a function of frequency. (C) Phase offset (absolute value) vs. distance (1-4 Hz). (D) Slope of phase offset (absolute value) vs. distance as a function of frequency (mean and 99% confidence intervals across sessions). Stars indicate significant differences before and during anesthetic effects (p&lt;0.01, corrected for multiple comparisons). Gray bar at 60 Hz due to line noise filtering effects.

Before drug delivery, Figure 4A (gray lines) shows that phase locking decreased with electrode distance within each array in the low-frequency range (1-4 Hz). Figure 4B (gray lines) shows that this effect existed across a wide range of frequencies (decrease indicated by all negative slopes). Phase offset with distance was positive but nearly flat before drug delivery for the low frequencies (gray lines, Fig. 4C). By contrast, the mid-range frequencies (15-40 Hz) showed marked increase in phase offset with distance before drug delivery (Fig. 4D), likely due to the greater relative power of mid-range frequencies during the awake state.

Both ketamine and dexmedetomidine amplified these trends. At low frequencies, the drugs caused a sharper decline in phase locking and sharper increase in phase offset with distance (Fig. 4A, C). Note that there are fewer pairs in an array at farther distances, which could lead to the greater variability shown, in addition to the decreased phase locking at those distances. At higher frequencies, the results were mixed. The effect of ketamine on phase locking in the gamma band (40-100 Hz) was similar to low frequencies (Fig. 4B). Dexmedetomidine, by contrast, caused a flattening of the slope of PLV vs. distance between 30-70 Hz. Neither drug had large effects on the overall phase offsets at high frequencies (Fig. 4D).

A continuation of these distance trends amplified under drug influence could explain the phase offsets between arrays in ventrolateral and dorsolateral PFC. The cortical surface distance between these arrays was ∼20 mm, so given the ∼20°-30° phase offset between channels at either end of the array (∼2.5 mm apart), extrapolating across distance could explain the observed ∼180° phase offset across regions. The phase locking at the farthest distance within an array is still above values seen between arrays, but likely levels off at further distances.

## Sub-anesthetic doses of ketamine and dexmedetomidine produced weaker effects

Sub-anesthetic doses of both drugs (1 mg/kg ketamine, 5 mg/kg dexmedetomidine) produced effects that were weak and more variable but trended towards those seen in anesthetic doses. For ketamine, there was no change in low-frequency power and a smaller increase in gamma power (Fig. 5A, left) in contrast to the anesthetic dose. For dexmedetomidine, there was a smaller increase in low-frequency power relative to the anesthetic dose (Fig 5A, right). Between arrays, there were significantly smaller effects in phase locking at low frequencies compared to the higher doses of each drug (horizontal bars) with few consistent trends within or across hemispheres (stars) (Fig. 5B). The phase offset effects were smaller under sub-anesthetic doses but often trended in the same direction (Fig. 5C-D), and they reached significance for fewer comparisons. Within the arrays, the local distance effects were reduced. Sub-anesthetic doses caused a smaller decrease in phase locking (Fig. 5E) and a smaller increase in phase offset (Fig. 5F) with distance than the anesthetic doses.

**Figure 5:**
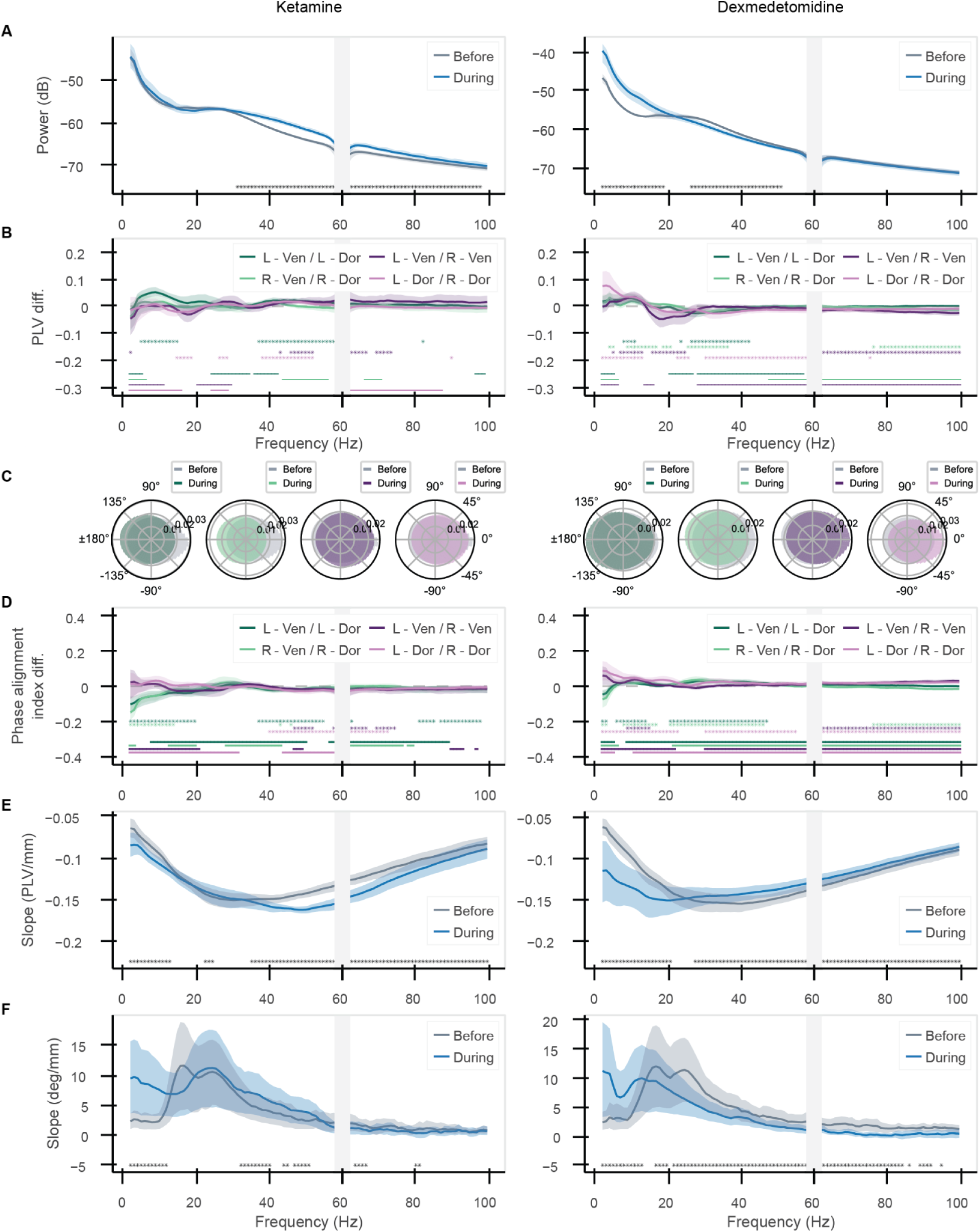
Sub-anesthetic doses of ketamine and dexmedetomidine produce weaker effects. (A) Average LFP power. (B) Change in PLV. (C) Histograms of phase offsets. (D) Change in phase alignment index. (E) Phase locking slope vs. frequency within array. (F) Phase offset (absolute value) slope vs. frequency within array. Lines and shading show mean and 99% confidence intervals across sessions. Stars indicate significant differences before and during anesthetic effects (p&lt;0.01, corrected for multiple comparisons). Colored bars in (B) and (D) indicate significant differences between low and high doses of anesthetics (p&lt;0.01, calculated with a permutation test, corrected for multiple comparisons). Gray bar at 60 Hz due to line noise filtering effects.

## Discussion

Ketamine and dexmedetomidine, while acting on different molecular and circuit pathways, induce similar changes in low-frequency oscillatory dynamics. These dynamics may be a marker of loss of responsiveness and suggest a fundamental role for broad, low-frequency activity in driving and disrupting consciousness.

We found that anesthetics heightened low-frequency cortical activity, misaligning this activity in different regions of PFC, while aligning it in homologous regions across hemispheres (depicted in Fig. 6A). The drugs had different effects across frequencies up to and including gamma, but the low frequency effects were the common feature between them. Increases in low-frequency power have been observed in cortical LFPs in animals and in electroencephalogram signals in humans for these drugs^13–18^ as well as propofol^19–24^, another anesthetic drug that is widely studied. We also noted increased phase locking of LFPs both within and across hemispheres, an effect that has previously been observed in various cortical regions with ketamine^17^, dexmedetomidine^16^, and propofol^23,24^.

**Figure 6:**
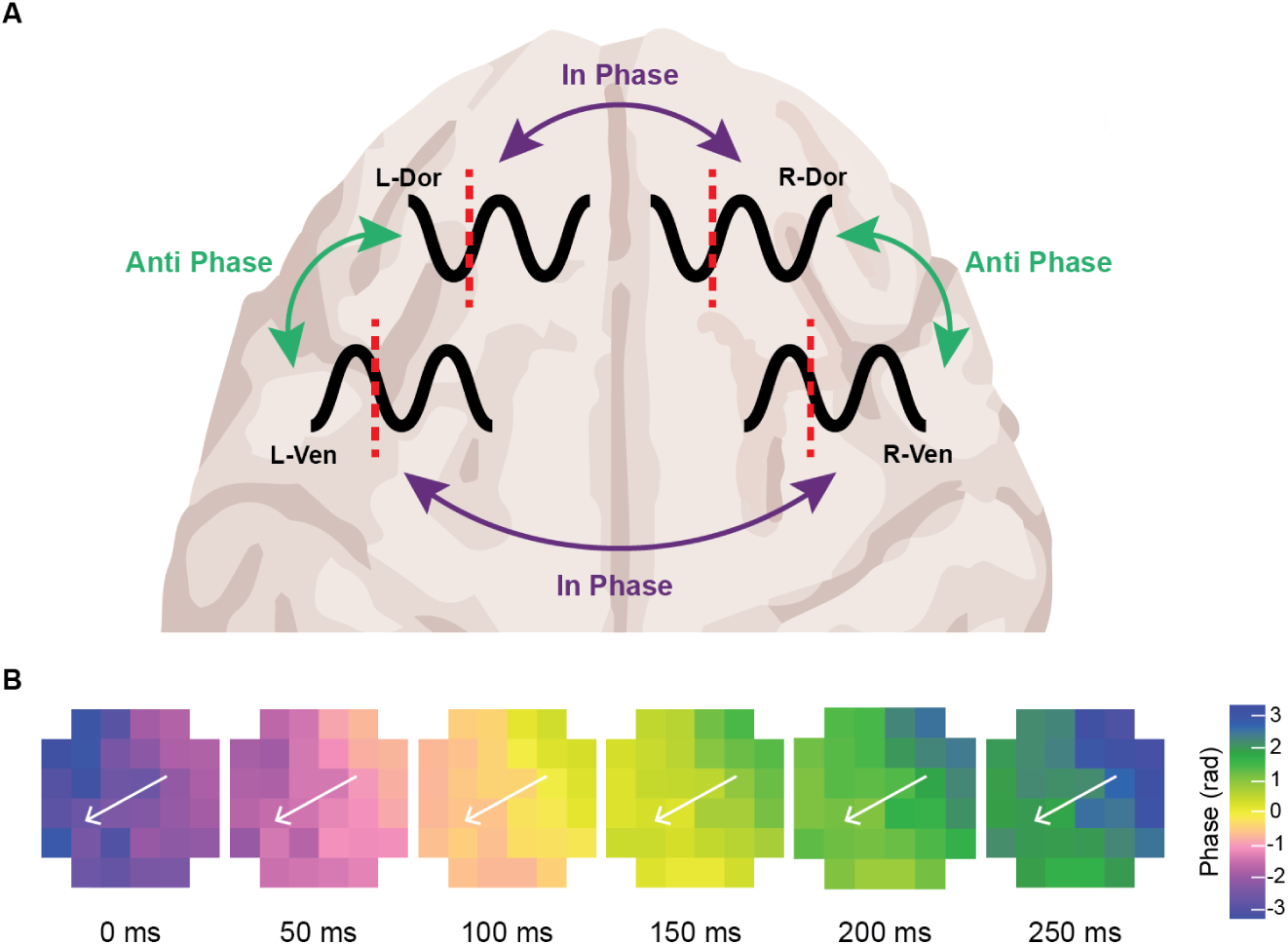
Anesthetics fragment cortical activity within a hemisphere, but synchronize it across hemispheres. (A) Ketamine and dexmedetomidine both cause phases of low-frequency oscillatory activity to become misaligned between different regions of PFC, and more aligned between homologous regions across hemispheres. (B) Example traveling wave. Phase of low frequency (1-4 Hz) LFP signal in an electrode array across time from a recording with ketamine. Each square represents one electrode contact, colored according to its instantaneous phase at the indicated time.

While past work has shown that both drugs increase phase locking across cortex at low frequencies, phase locking can be caused by different underlying patterns of neural activity, and such activity need not be the same across different anesthetic drugs. Here, we find specific patterns of phase alignment that are consistent across the two drugs. Within a hemisphere, the ventrolateral and dorsolateral PFC became more anti-phase at low frequencies, with their phase offsets trending toward 180°. The low-frequency offsets that we observed were consistent across entire arrays, suggesting the observed effects come from spatially broad oscillations.

Additionally, we found that extrapolating local phase-offset effects within a region across longer distances could explain the anesthetic-induced 180° phase offset between the ventrolateral and dorsolateral PFC. These effects could be due to a large traveling wave that moves between the areas (Fig. 6B shows an example from our recordings of such a wave traveling across a single electrode array with anesthesia).

Large-scale cortical traveling waves are known to occur spontaneously during anesthesia^25–27^. These waves may play an essential role in shaping neural activity when awake^28–32^, and severely limiting it during anesthesia^27,33^ by fragmenting within-hemisphere communication as periods of excitability become misaligned^3–8,34^. In fact, propofol has been shown to fragment activity and decrease information transfer across long distances (>2 cm) in the human cortex^35–37^, and to prevent sensory signals from reaching the macaque frontal cortex^38^. We note that there was not a complete misalignment of phases at all times, but rather a tendency for LFPs to be out of phase more often. Such “relative coordination” can still strongly influence network activity^39^.

In contrast to the observed fragmentation of activity within a hemisphere, we found that homologous regions across hemispheres became more aligned. The increase in interhemispheric alignment of activity by anesthetics seems to reverse the functional distinctions observed in the awake, cognitively-engaged brain. In that case, the two hemispheres are often able to function semi-independently, at least for visual cognition^40^. During quiet awake states and anesthesia, however, cross-hemispheric synchrony increases in mice^41^, in line with our observations. This synchrony was found to be dependent upon the presence of the intact corpus callosum^41^, which is known to preferentially connect homologous regions^42^. Such synchrony could also be mediated by subcortical inputs, such as those from the thalamus, a known hub of interhemispheric communication^43,44^.

It is possible that our observations of low-frequency changes in LFP signals can be explained by the activity of cortical layer 5 pyramidal neurons, which anesthetics are known to influence^45–49^.

These neurons make large contributions to the LFP and have low pass filtering properties at their somas^50–52^. Thus, they may play a significant role in the increases in power and phase locking of cortical slow-wave dynamics.

These neurons are also known to be a prominent component of thalamocortical loops^53^. The molecular effects of these drugs may cause loss of responsiveness through common disruptions to thalamocortical interactions^54,55^, via both cortical and subcortical action. Thalamocortical connectivity is hypothesized to play a major role in consciousness^7,47,56–60^ and in anesthetic-induced dissociation^61^ or unconsciousness^23,24,62–68^. Experimental and modeling work show that low-frequency oscillations and thalamocortical connections may be involved in the transfer of information between cortical regions^69,70^, and that such transfer may be disrupted during anesthesia, thus leading to unconsciousness^38,67,68^. Simultaneous recordings from and stimulation of the cortex and thalamus during anesthesia will help to illuminate the role that these interactions play in driving the broad phase-offset activity that we found here.

Our results suggest that the loss of responsiveness caused by anesthetic doses of ketamine and dexmedetomidine is associated with changes not only in oscillatory power, but also in the alignment of low-frequency signals within and between cerebral hemispheres. These effects were severely weakened by sub-anesthetic doses of the drugs that were not sufficient to cause loss of responsiveness. Despite their different mechanisms of action at the molecular level, we found similar network-level effects between the two drugs. Thus, cortical network-level activity may be the optimal lens through which we can understand the convergent mechanisms of different anesthetics, and even make use of this understanding to inform anesthetic administration^71^, as well as mechanisms of^72–76^ and treatments for^77–79^ psychiatric diseases.

## Methods

### Experimental design

All surgical and animal care procedures were approved by the Massachusetts Institute of Technology (MIT)’s Committee on Animal Care and were conducted in accordance with the guidelines of the National Institute of Health and MIT’s Division of Comparative Medicine. Ketamine and dexmedetomidine were administered as an intramuscular bolus to two monkeys (Monkey S and Monkey P, both males) at sub-anesthetic (1 mg/kg ketamine, 5 mg/kg dexmedetomidine) and anesthetic (10 mg/kg ketamine, 20 mg/kg dexmedetomidine) doses.

There were 13 subanesthetic dose ketamine sessions (6 with Monkey P, 7 with Monkey S), 16 subanesthetic dose dexmedetomidine sessions (8 with Monkey P, 8 with Monkey S), 12 anesthetic dose ketamine sessions (6 with Monkey P, 6 with Monkey S), and 16 anesthetic dose dexmedetomidine sessions (8 with Monkey P, 8 with Monkey S). One anesthetic ketamine session with monkey P was excluded from analysis due to noise from an unknown source.

Induction of anesthetic effects was measured by loss of responsiveness in a lever-pressing task. In this task monkeys were trained to press a lever in response to a tone and received a juice reward for a correct response. All monkeys responded regularly before anesthetic injection, then stopped responding to the lever-pressing task before the period from which the “during anesthetic” data was taken for analysis. Changes in spectral LFP power consistent with known effects of ketamine and dexmedetomidine began around 3-4 minutes after anesthetics were injected (Fig. 1B) and could be seen throughout the period analyzed during anesthetics (Fig. 1C), which began 10 minutes after drug administration.

To study the effects of anesthetics on neural synchrony in PFC, neural activity was recorded continuously throughout the experiment (before, during, and after anesthetic injection) with chronically implanted Utah arrays placed in the left and right ventrolateral PFC and dorsolateral PFC (referred to as L-Ven, R-Ven, L-Dor, and R-Dor). Arrays were placed on either side of the principal sulcus but varied slightly in exact location and orientation. Each array had a regularly spaced grid of 6x6 electrodes, 400 micrometers apart (minus the four corner locations, for a total of 32 channels per array).

In periods before and during anesthetics, the monkeys were presented with a separate passive auditory task, in which varying tones were played but monkeys were not required to respond. All analysis was performed on time periods during which the auditory task was happening. Results were indistinguishable during and between blocks of sounds within the auditory task period, so we pooled data across the entire period for analysis.

### LFP processing

LFPs were recorded at 1kHz. All channels were referenced to a wire inserted under the dura in one of the craniotomies. For sessions with Monkey P (excluding three anesthetic-dose ketamine sessions), noise was removed at 60, 120, and 180 Hz with online Butterworth notch filtering, resulting in a dip in power around 60 Hz (higher frequencies were not analyzed here). For Monkey S (and the three monkey P sessions), filtering was performed offline, after data collection. We estimated the line noise at the 1st through 8th harmonics of 60 Hz (i.e., 60-480 Hz in steps of 60) by fitting the raw data on each channel with a temporally adaptive sine function at each given frequency. The fitted sinusoids were then subtracted from the data. Due to the effects of the notch filtering, the 60 Hz region is blocked off in further analyses.

To remove common noise and signal from the reference in the LFPs, we used Lepage et al.’s robust common average reference (rCAR)^9^. Many methods exist for removing noise from LFPs, which each have different benefits and drawbacks^80,81^; we chose this method due to its reduced impact on phase relationships. Methods such as no re-referencing or global mean subtraction both distort phase relationships between signals. No re-referencing leads to an increase in phase alignment at the frequency of any common noise or signal inserted by the reference channel. Global mean subtraction can induce spurious phase offsets^82^. rCAR uses a robust maximum-likelihood type estimator to calculate the reference signal, and is shown to be empirically better at preserving the phase of signals than a global mean reference^9^. We compare these methods in Figure S6, finding similar trends in the changes in phase relationships during anesthesia, but slight variation in the overall phase distributions for each method.

To identify outlier channels, we calculated the standard deviation of the LFP from each electrode in a session. We then found the interquartile range of standard deviations for all channels in a session, and we removed channels with a standard deviation more than 5 interquartile ranges outside of the overall range, with an average of 3.8 channels out of 128 removed per session.

### Spectral analysis

We used multitaper spectral estimation on a series of overlapping time windows to analyze data in the frequency domain. LFPs were split into 1 second time windows, beginning every 0.1 seconds (so each window had 0.9 seconds overlap with the next one). We selected 1 second time windows because ketamine anesthesia causes up and down states in which alternating bands of frequency are higher in power, and these states have been found to last about 1 second on the lower end in monkeys^18^. However, we found that the results did not change significantly when using a 2 second time window (Fig. S6). We calculated the multitaper spectrogram^83–85^ for each window using a time-half bandwidth product of 2 with 3 tapers (as in ^18^) over a total frequency range of 1-100 Hz.

### PLV, coherence, phase offset

PLV, coherence, and phase offset were calculated between pairs of channels on indicated arrays. For a comparison between two given arrays, we analyzed all possible pairs of channels between the two arrays. We calculated the cross spectrum C_XY_ by multiplying signal 𝑋 by the complex conjugate 𝑌*of signal 𝑌 (in the frequency domain) at each time window/taper for each channel pair. We also calculated auto spectra C_XY_ and C_XY_ by multiplying a signal by its own complex conjugate at each time window/taper.

To calculate the PLV, we took the absolute value of the average normalized cross spectrum. To calculate coherence, we divided the cross spectrum of two channels by the square root of the product of each channel’s auto spectrum. To calculate the phase offset, we took the angle of the cross spectrum. We then averaged over the tapers. For within-array analyses, we took all possible pairings of channels within an array (excluding self-pairs) and then sorted the pairings by distance between sites.

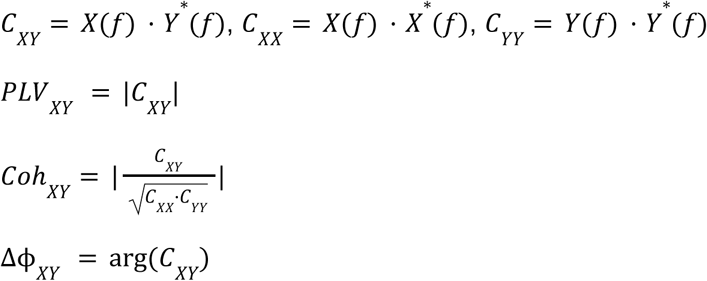

PLV and coherence are dimensionless measures ranging from 0 to 1. Phase offsets range from -180° to 180°. All line plots show mean values with 99% confidence intervals. We first averaged across all channels or channel pairings for the indicated regions within a session. We then computed averages and nonparametric bootstrap confidence intervals across sessions (100,000 bootstrap replicates).

### Phase histograms

Phase offset histograms show the distribution of phase offsets between all channel pairs in indicated regions, accumulated across all time points within a given recording period (before or during anesthetics). Phase offsets were calculated as indicated above and normalized by the total count of phase values contained in each histogram. The histograms show the distribution over time points and channel pairs (neither time nor channel pairs were averaged out). The histograms have 50 bins, so a uniform distribution across all phase offsets would result in values of 0.02 across all bins. We tested the histograms with finer-grained binning (500 bins) to ensure that no aspects of the distribution were occluded by the choice of bin size and confirmed that they were qualitatively similar. The histograms show phase offsets for the 1-4 Hz frequency range.

Phase alignment index (as in Fig. 3C-E) was calculated by taking the cosine of each phase offset and averaging over time and channel pairs within a session. We chose to take the cosine of the phase offset because the peaks of the phase histograms tended towards 0° and 180°. In this metric, a circular uniform distribution of phases has an average value of zero, while distributions with more weight towards 0° or 180° have positive or negative values, respectively. Note that distributions weighted towards +/-90° would also have a phase alignment index of zero; however, the distributions investigated here were not weighted in those directions. Phase alignment index is a dimensionless measure ranging from -1 to 1.

### Instantaneous phase

Instantaneous phase of the example traveling wave in Figure 6 was calculated by band passing the LFP using a 3rd order Butterworth filter in the 1-4 Hz range then taking the Hilbert transform and calculating the angle of the signal at the indicated time points.

## Statistics

Line plots represent averages of session averages and shaded regions represent 99% confidence intervals on across-session averages. Confidence intervals are nonparametric bootstraps across session means with 100,000 replicates. Stars indicate significant differences between metrics before and during anesthesia at p<0.01 level, calculated with a pairwise permutation test using 100,000 replicates. Bars (in Figure 5B, D) indicate significant differences between metrics calculated for low and high doses of anesthetics at p<0.01 level, calculated with a permutation test using 100,000 replicates. P-values were corrected for multiple comparisons across the different frequencies using the Benjamini/Hochberg procedure for controlling the false discovery rate.

## Acknowledgements

The authors thank Sebastian Gallo and the MIT Division of Comparative Medicine for assistance with monkey care. This material is based upon work supported by the

U.S. Department of Energy, Office of Science, Office of Advanced Scientific Computing Research, under Award Number DE-SC0024386 (A.G.B.). This work was also funded by NIMH 1R01MH131715-01 (E.K.M.), NIMH R01MH11559 (E.K.M.), NIH 1R21AG077275-01A1 (Y.I.), The Simons Center for the Social Brain (E.K.M.), The JPB Foundation (E.N.B., E.K.M.), and The Picower Institute for Learning and Memory (E.N.B., E.K.M.).

## Disclaimer

This report was prepared as an account of work sponsored by an agency of the United States Government. Neither the United States Government nor any agency thereof, nor any of their employees, makes any warranty, express or implied, or assumes any legal liability or responsibility for the accuracy, completeness, or usefulness of any information, apparatus, product, or process disclosed, or represents that its use would not infringe privately owned rights. Reference herein to any specific commercial product, process, or service by trade name, trademark, manufacturer, or otherwise does not necessarily constitute or imply its endorsement, recommendation, or favoring by the United States Government or any agency thereof. The views and opinions of authors expressed herein do not necessarily state or reflect those of the United States Government or any agency thereof.

## Author contributions

Conceptualization, A.G.B., J.J.B., S.L.B., Y.I., E.N.B., E.K.M.;

Investigation J.J.B., J.E.R., M.K.M.; Data curation, J.J.B., S.L.B.; Formal Analysis, A.G.B.,

S.L.B., E.K.M.; Supervision, Y.I., E.N.B, E.K.M; Writing – original draft, A.G.B, E.K.M.; Writing –

review & editing, A.G.B., J.J.B., S.L.B., E.K.M.; Funding acquisition, E.N.B., E.K.M.

---

**Figure S1:**
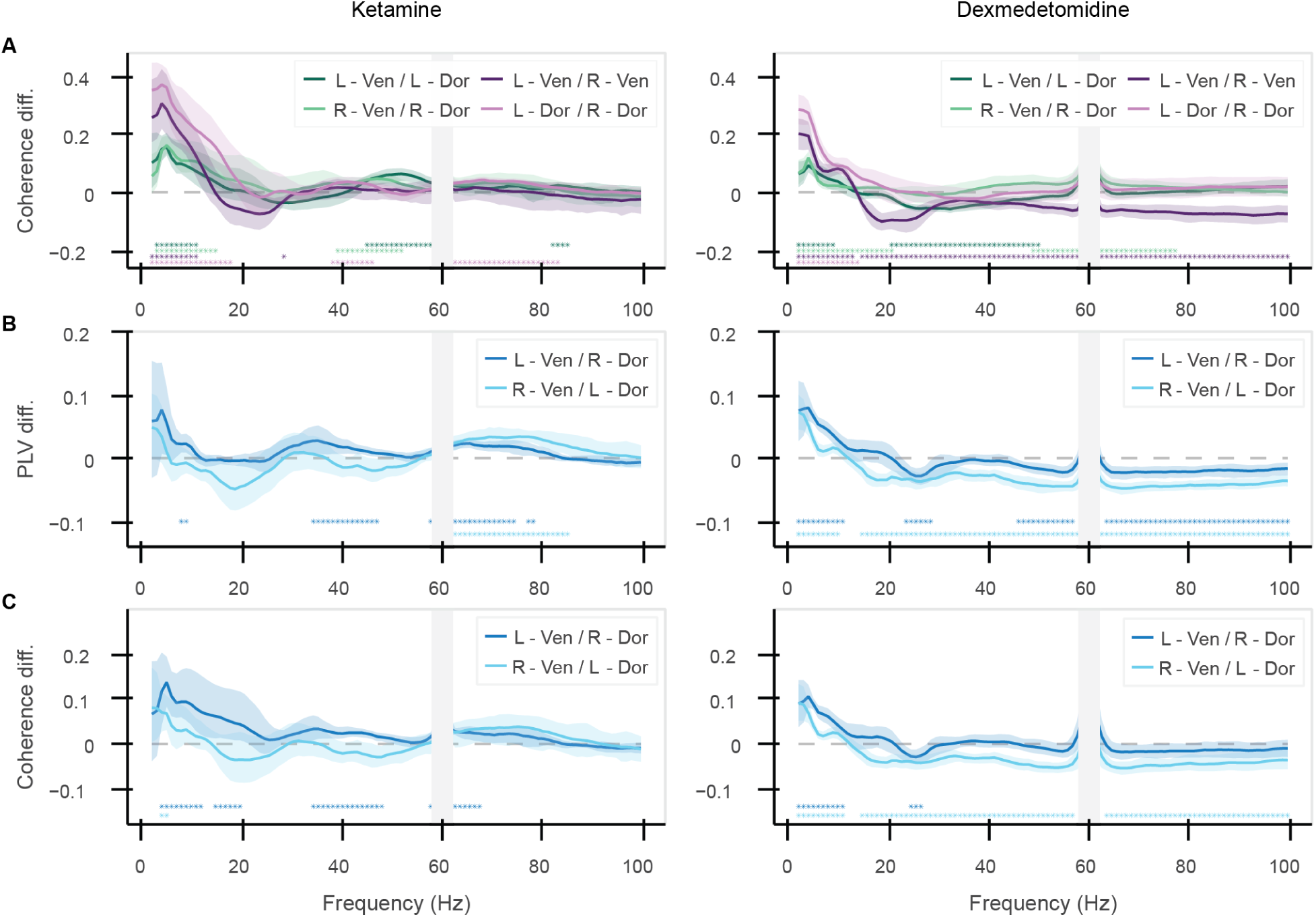
Results were similar for coherence measure and for cross-area interhemispheric comparisons. (A) Coherence within and across hemispheres. (B) PLV for cross-hemisphere / cross-region pairs. (C) Coherence for cross-hemisphere / cross-region pairs. All plots show differences (during – before anesthetics) (mean and 99% confidence intervals across sessions). Stars indicate significant differences before and during anesthetic effects (p&lt;0.01, corrected for multiple comparisons). Gray bar at 60 Hz due to line noise filtering effects.

**Figure S2:**
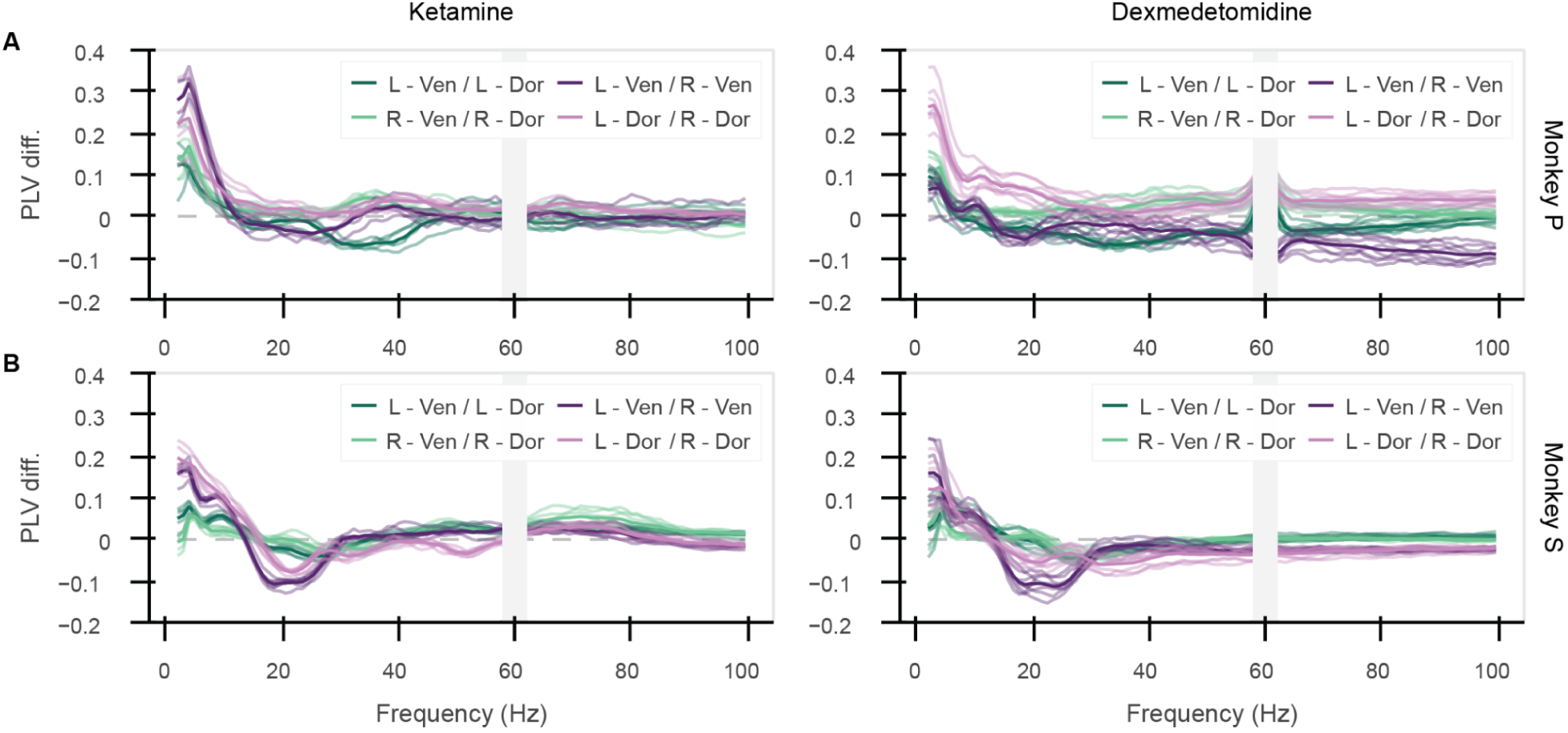
Changes in phase locking were consistent across monkeys, sessions. Difference in PLV (during – before anesthetics) for two monkeys (A) Monkey P and (B) Monkey S. Each line indicates one session; the darker line is the average across all sessions. Gray bar at 60 Hz due to line noise filtering effects.

**Figure S3:**
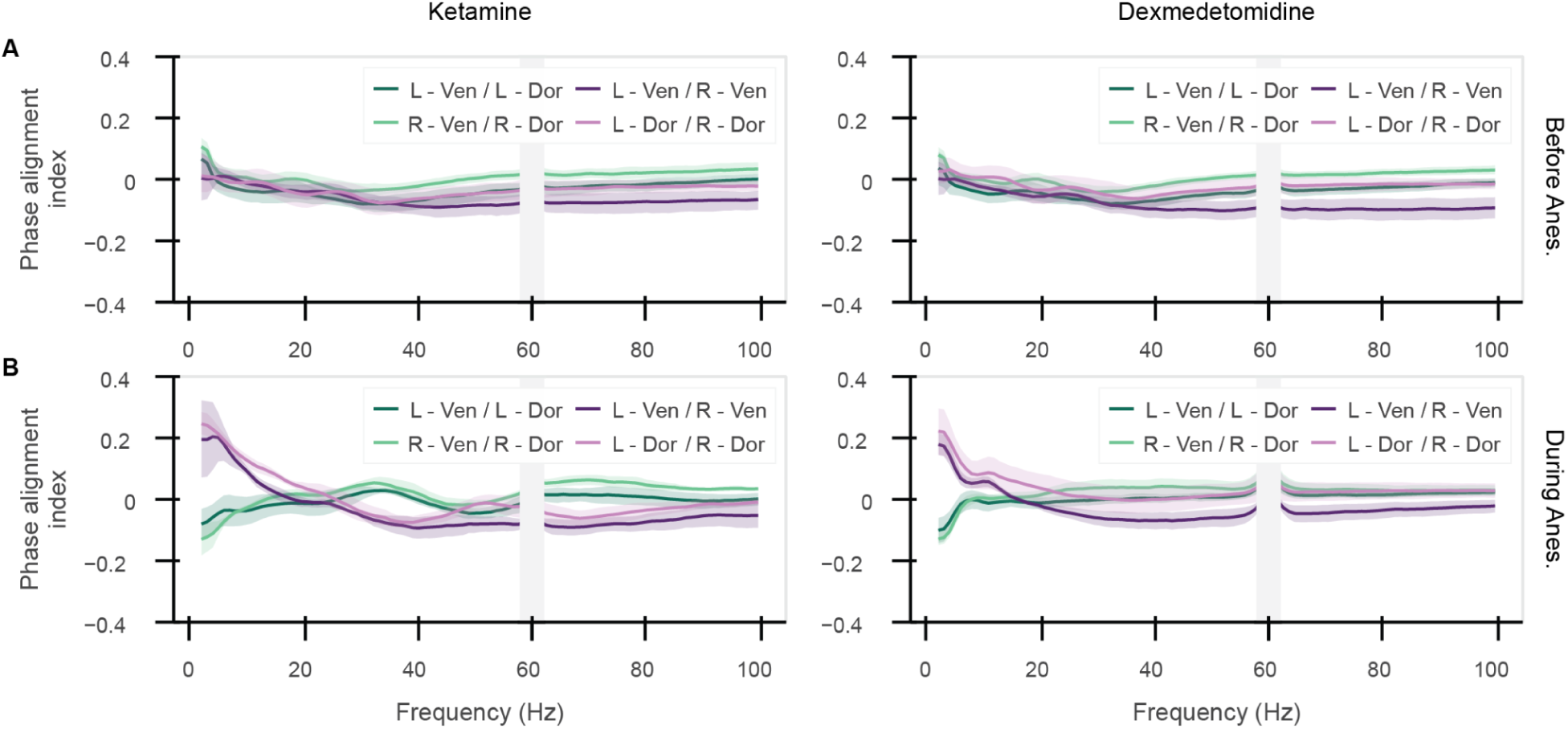
Phase alignment shifted with anesthetics. Phase alignment index cos(Δϕ) (A) before anesthetics, and (B) during anesthetics (mean and 99% confidence intervals across sessions). Difference between (A) and (B) shown in Figure 3C. Gray bar at 60 Hz due to line noise filtering effects.

**Figure S4:**
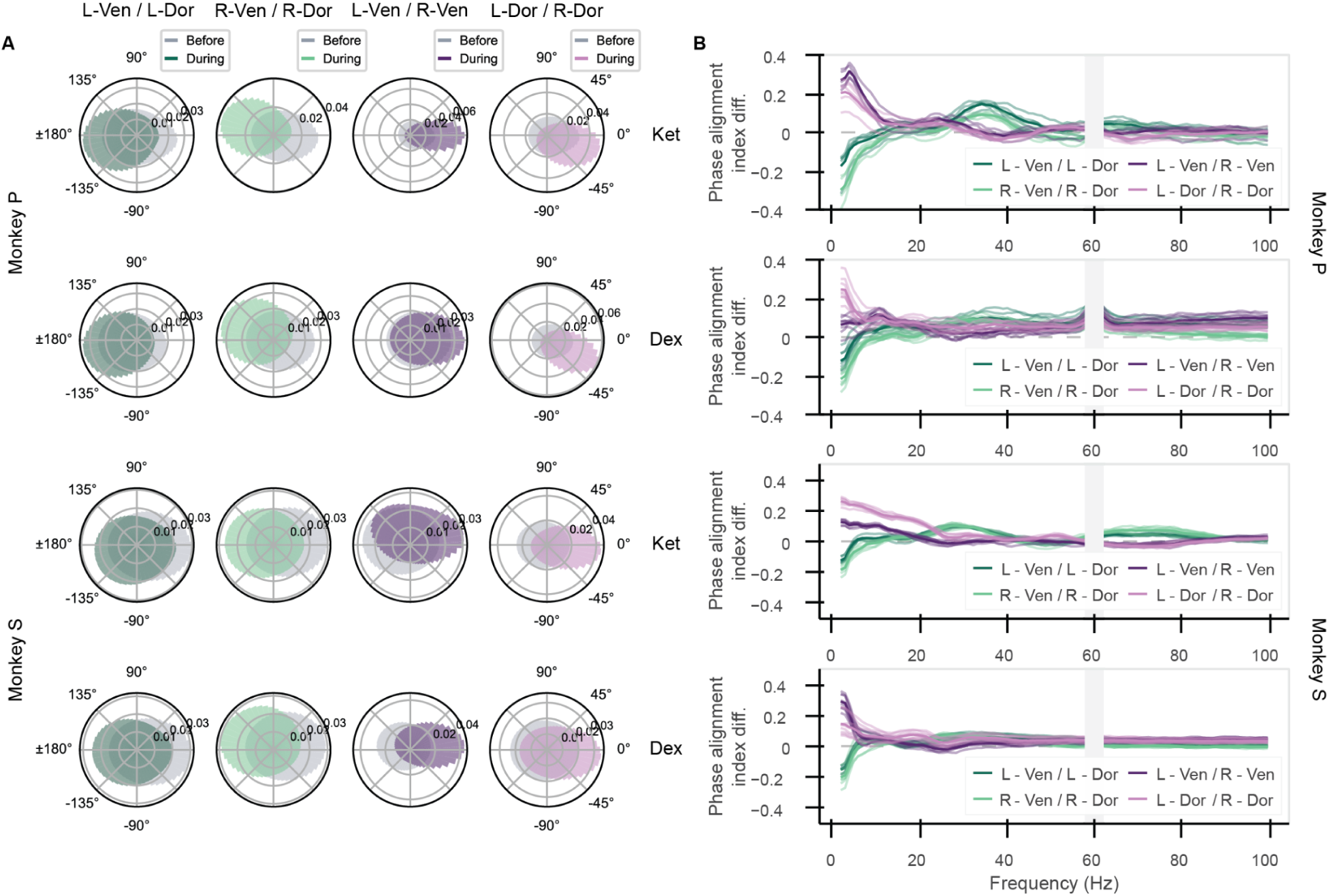
Shift in phase alignment was consistent across monkeys, sessions. (A) Histograms of phase offsets in the 1-4 Hz range between indicated regions for each drug, split by monkey (top two rows are Monkey P, bottom two rows are Monkey S). (B) Changes in phase alignment index for each drug and monkey (corresponding to rows in A). Each line indicates one session; the darker line is the average across all sessions. Gray bar at 60 Hz due to line noise filtering effects.

**Figure S5:**
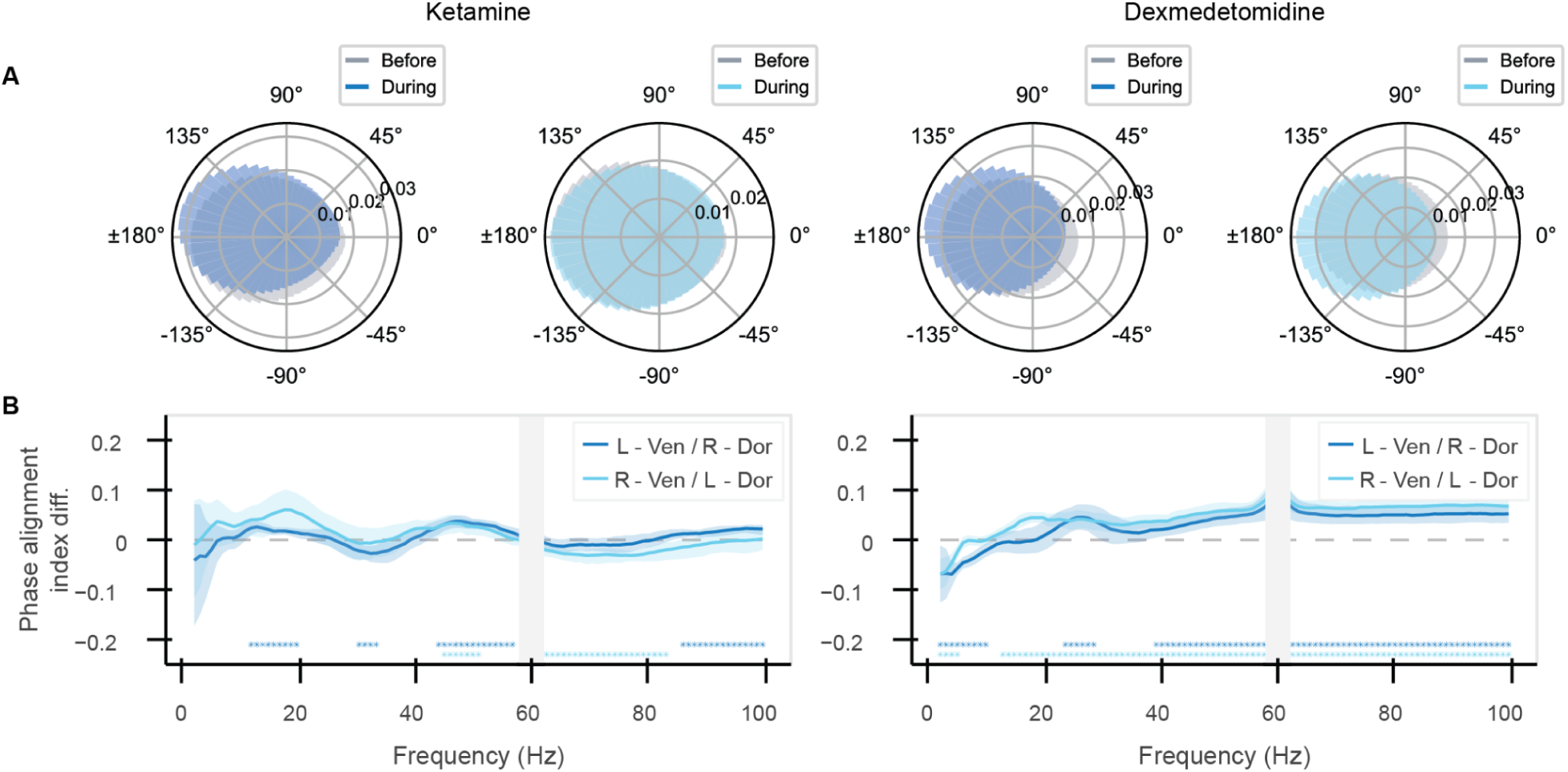
Phase alignment decreased slightly between diagonal regions at low frequencies. (A) Phase offsets of cross-hemisphere / cross-region pairs. (B) Changes in cross-hemisphere / cross-region phase alignment index (mean and 99% confidence intervals across sessions). Stars indicate significant differences in phase alignment before and during anesthetic effects (p&lt;0.01, corrected for multiple comparisons). Gray bar at 60 Hz due to line noise filtering effects.

**Figure S6:**
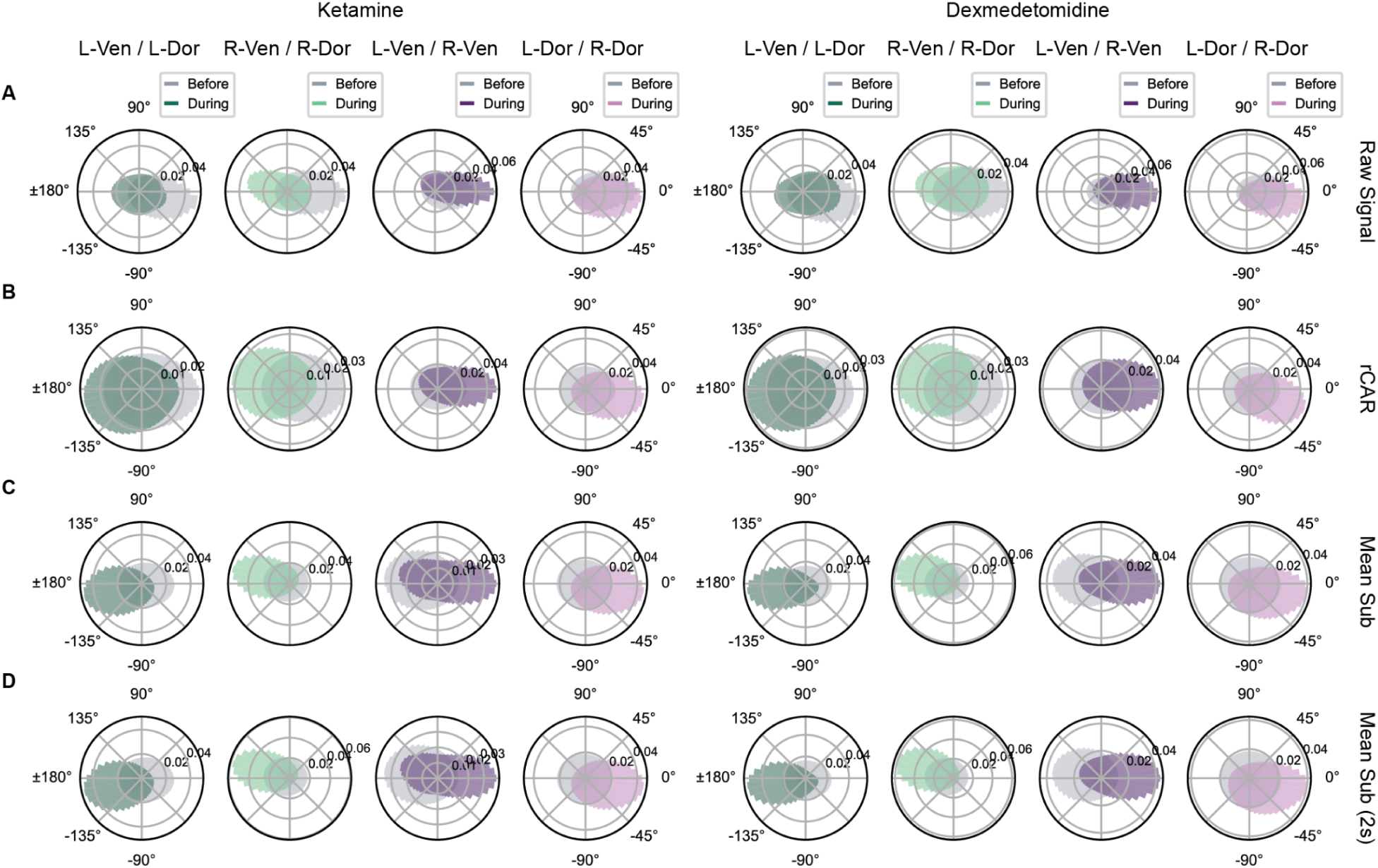
Change in phase alignment followed the same trend with different LFP referencing techniques. (A) Phase offsets calculated with raw LFP signal. Offsets are more in-phase compared to other techniques. (B) Phase offsets calculated with robust common average reference technique. (C) Phase offsets calculated with global mean subtraction. Offsets are more anti-phase compared to other techniques. (D) Phase offsets calculated with global mean subtraction and 2 second spectral time window. Offsets are similar to 1 second time window with the same technique (C).

